# Minimum Information for Reporting Next Generation Sequence Genotyping (MIRING): Guidelines for Reporting HLA and KIR Genotyping via Next Generation Sequencing

**DOI:** 10.1101/015230

**Authors:** Steven J. Mack, Robert P Milius, Benjamin D Gifford, Jürgen Sauter, Jan Hofmann, Kazutoyo Osoegawa, James Robinson, Mathjis Groeneweg, Gregory S Turenchalk, Alex Adai, Cherie Holcomb, Erik H Rozemuller, Maarten T Penning, Michael L Heuer, Chunlin Wang, Marc L Salit, Alexander H Schmidt, Peter R Parham, Carlheinz Müller, Tim Hague, Gottfried Fischer, Marcelo Fernandez-Viňa, Jill A Hollenbach, Paul J Norman, Martin Maiers

## Abstract

The development of next-generation sequencing (NGS) technologies for *HLA* and *KIR* genotyping is rapidly advancing knowledge of genetic variation of these highly polymorphic loci. NGS genotyping is poised to replace older methods for clinical use, but standard methods for reporting and exchanging these new, high quality genotype data are needed. The Immunogenomic NGS Consortium, a broad collaboration of histocompatibility and immunogenetics clinicians, researchers, instrument manufacturers and software developers, has developed the Minimum Information for Reporting Immunogenomic NGS Genotyping (MIRING) reporting guidelines. MIRING is a checklist that specifies the content of NGS genotyping results as well as a set of messaging guidelines for reporting the results. A MIRING message includes five categories of structured information – message annotation, reference context, full genotype, consensus sequence and novel polymorphism – and references to three categories of accessory information – NGS platform documentation, read processing documentation and primary data. These eight categories of information ensure the long-term portability and broad application of this NGS data for all current histocompatibility and immunogenetics use cases. In addition, MIRING can be extended to allow the reporting of genotype data generated using pre-NGS technologies. Because genotyping results reported using MIRING are easily updated in accordance with reference and nomenclature databases, MIRING represents a bold departure from previous methods of reporting *HLA* and *KIR* genotyping results, which have provided static and less-portable data. More information about MIRING can be found online at miring.immunogenomics.org.

---

**Abbreviations**
BAM: – Binary Alignment/Map
CSB: – Consensus Sequence Block
dbGAP: – Genotype and Phenotype Database
EMBL: – European Molecular Biology Laboratory
ENA: – European Nucleotide Archive
GL: – Genotype List
GRC: – Genome Reference Consortium
GTR: – Genetic Testing Registry
HLA: – Human Leukocyte Antigen
HIEDFS: – HLA Information Exchange Data Format Standards
HIPAA: – Health Insurance Portability and Accountability Act
IEC: – International Electrotechnical Commission
IDAWG: – Immunogenomic Data Analysis Working Group
IHIW: – International HLA and Immunogenetics Workshop
IMGT: – ImMunoGeneTics
INGSDC: – Immunogenomic Next Generation Sequencing Data Consortium
INSDC: – International Nucleotide Sequence Database Collaboration
IPD: – Immuno Polymorphism Database
ISO: – International Organization for Standardization
IUBMB: – International Union of Biochemistry and Molecular Biology
IUPAC: – International Union of Pure and Applied Chemistry
KIR: – Killer-cell Immunoglobulin-like Receptor
MIBBI: – Minimum Information for Biological and Biomedical Investigations
MIRING: – Minimum Information for Reporting Immunogenomic NGS Genotypes
NCBI: – National Center for Biotechnology Information
NGS: – Next Generation Sequencing
OID: – Organization Identifier
PIPEDA: – Personal Information Protection and Electronics Documents Act
SBT: – Sanger sequencing Based Typing
SFF: – Standard Flowgram Format
SRA: – Sequence Read Archive
SSOP: – Sequence-Specific Oligonucleotide Probe
SSP: – Sequence-Specific Priming
URI: – Uniform Resource Identifier
VCF: – Variant Call Format

## 1. Introduction

Next-generation sequencing (NGS) offers high-throughput generation of phased sequences for the highly polymorphic human leucocyte antigen (*HLA*) and killer-cell immunoglobulin-like receptor (*KIR*) genes, allowing their rapid, high-resolution genotyping. NGS methods may be more generally described as single-molecule sequencing methods[1]. In some cases, these methods offer full-gene sequence results[1–4]. In general, all NGS methods offer higher resolution and lower ambiguity genotypes than standard methods such as “Sanger” sequencing based typing (SBT), and sequence-specific oligonucleotide probe (SSOP) or primer (SSP) methods[3–6], and do not require the use of secondary genotyping methods to resolve ambiguities.

Any method for genotyping *HLA* and *KIR* using genomic DNA requires at least three components: the *genotyping instrument, reference sequences,* and *analysis software.* The *genotyping instrument* generates primary sequence data, which is interpreted by the *analysis software,* using the *reference sequences* to identify the subject’s genotype. A wide variety of instruments, reference sequence resources, and data analysis programs are available for both NGS and pre-NGS genotyping approaches, and are used in different combinations.

In some cases, the different methods may not generate the same results for a given subject. Such discrepancies may derive from the instrumentation, reference sequences, software, or a combination of these components. However, as Hollenbach et al.[7] have described, there is no standard format for reporting a genotyping result or for documenting the components that were applied to generate that result. In the absence of such documentation, the source of discrepancies in genotyping results is rarely identifiable. In addition, it becomes impossible to directly relate the *HLA* and *KIR* genotypes of subjects genotyped using different methods, as genetic differences between individuals may not be distinguishable from methodological differences between genotyping approaches. This lack of clarity has important implications for meta-analytical approaches to population or disease association studies that seek to combine and compare data across different studies. In general, ambiguity regarding the source of genotyping discrepancies impedes technical advances and optimization, and frustrates reproducible research.

Guidelines for reporting and documenting genotyping results are essential for evaluating *HLA* and *KIR* genotypes generated using different instruments, reference sequences or data-analysis programs. The active and ongoing development of NGS methods requires the adoption of a single extensible and adaptable standard for reporting and documenting NGS genotyping results.

Here we describe the Minimum Information for Reporting NGS Genotyping (MIRING) checklist, a set of Minimum Information for Biological and Biomedical Investigations (MIBBI)[8, 9] reporting guidelines developed by a consortium of immunogenomic researchers and clinicians, NGS instrument manufacturers and software developers, *HLA* and *KIR* sequence database developers and administrators, bone marrow donor registries and donor centers.

## 2. Description of MIRING

### 2.1 MIRING Development

The standard reporting of *HLA* and *KIR* genotypes is a long unmet need of the histocompatibility and immunogenetics community [7, 10–14]. The specific need for NGS genotype reporting guidelines emerged from a survey of Immunogenomic data management and analysis practices[15], carried out by the Immunogenomic Data Analysis Working Group (IDAWG) as part of the 16^th^ International HLA and Immunogenetics Workshop (IHIW)[16]. The survey uncovered a lack of consistency between laboratories and the resulting impact on downstream analytical results. The development of MIRING began with the formation of the Immunogenomic Next Generation Sequencing Data Consortium (INGSDC) (ngs.immunogenomics.org) by the IDAWG and the HLA Information Exchange Data Format Standards (HIEDFS) group. The INGSDC met several times between 2012 and 2014, and identified the minimum information needed to accurately report NGS genotyping results for the *HLA* and *KIR* genes for clinical and research applications. Further MIRING development took place as part of the Be The Match Foundation’s Data Standards ‘Hackathons’ for NGS-based typing held in September of 2014 and February of 2015 (dash.immunogenomics.org). Implementations of MIRING are being evaluated as part of a 17^th^ IHIW Informatics Component (ihiws.org/informatics-of-genomic-data/) project; bioinformatic tools for generating, exchanging and consuming MIRING messages are being developed as part of this project as well. The participation of interested investigators in this IHIWS project is welcome.

### 2.2 MIRING Goals

To meet the current needs of the histocompatibility and immunogenetics community for reporting and exchanging NGS genotype data, the elements of a MIRING message were designed with the following goals:

1. To facilitate downstream analyses and data management for current research and clinical use cases for molecular genotyping data in the histocompatibility and immunogenetics field.
2. To permit the re-analysis of NGS *HLA* or *KIR* genotyping results in the context of past, present and (foreseeable) future molecular nomenclatures and methods of describing HLA and KIR allele diversity.
3. To permit the comparison and evaluation of genotyping performance between different NGS platforms and analysis methods.
4. To enable molecular genotyping results generated using SBT, SSOP and SSP genotyping technologies to be incorporated if required.
5. That the MIRING elements be sufficient to permit the accurate reporting of NGS data generated for other highly-polymorphic regions of the human genome.

### 2.3. MIRING Elements

MIRING is both a checklist of elements that constitute a NGS *HLA* or *KIR* genotyping result, and a set of messaging guidelines for transmitting that NGS *HLA* or *KIR* genotyping result. Genotyping reports can be generated from a MIRING message. The MIRING guidelines include semantic definitions for a MIRING message, but are not intended to impose syntactic constraints on the message; they are principles that must be met, regardless of the structure of the message.

MIRING comprises eight primary elements, and their constituents (Table 1). Elements 1–5 constitute the MIRING message, suitable for reporting a genotyping result. Elements 6–8 constitute the contextual resource for MIRING messages, but are not included in MIRING messages; instead, these elements are referenced in MIRING messages. Where possible, MIRING elements are consistent with established formats for describing genetic and genomic data (e.g., FASTA[17–19], FASTQ[20], Variant Call Format (VCF)[21] and Genotype List (GL) String formats[22]), and leverage existing genetic and genomic data-resources (e.g., the IMGT/HLA and IPD-KIR Databases[23], the NCBI Genetic Testing Registry (GTR)[24] and International Nucleotide Sequence Database Collaboration (INSDC)[25]).

**Table 1.**
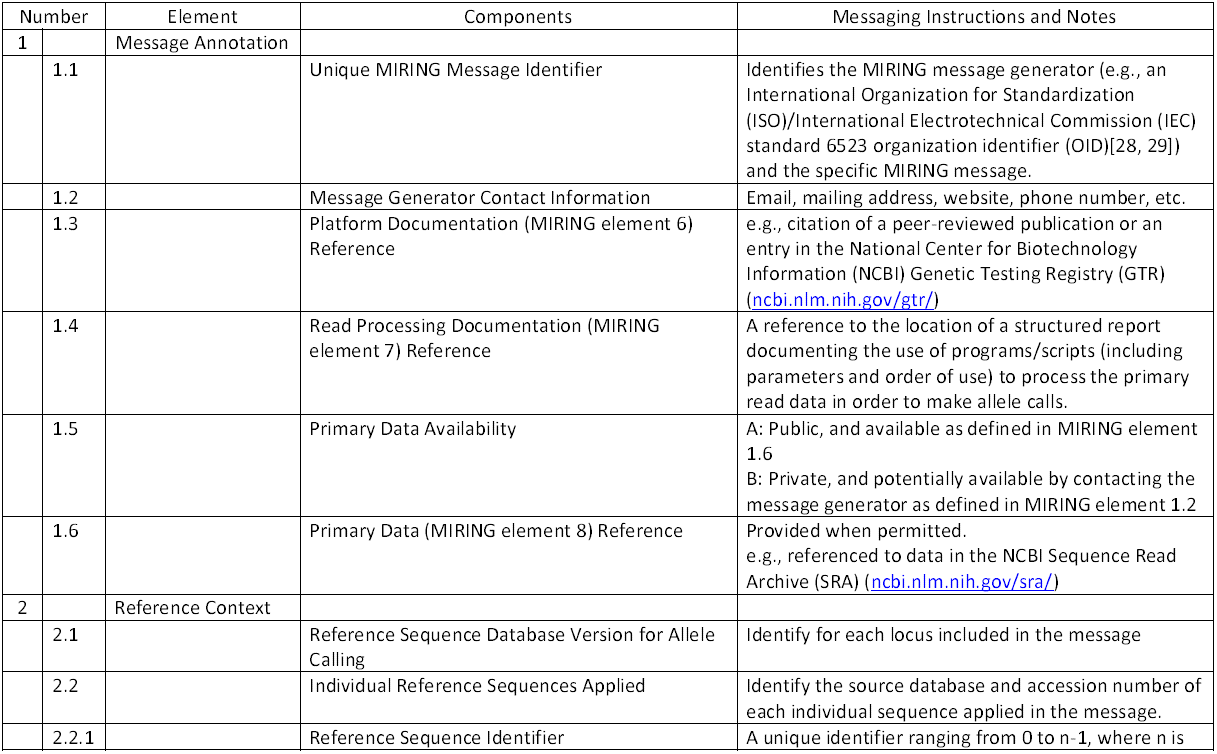
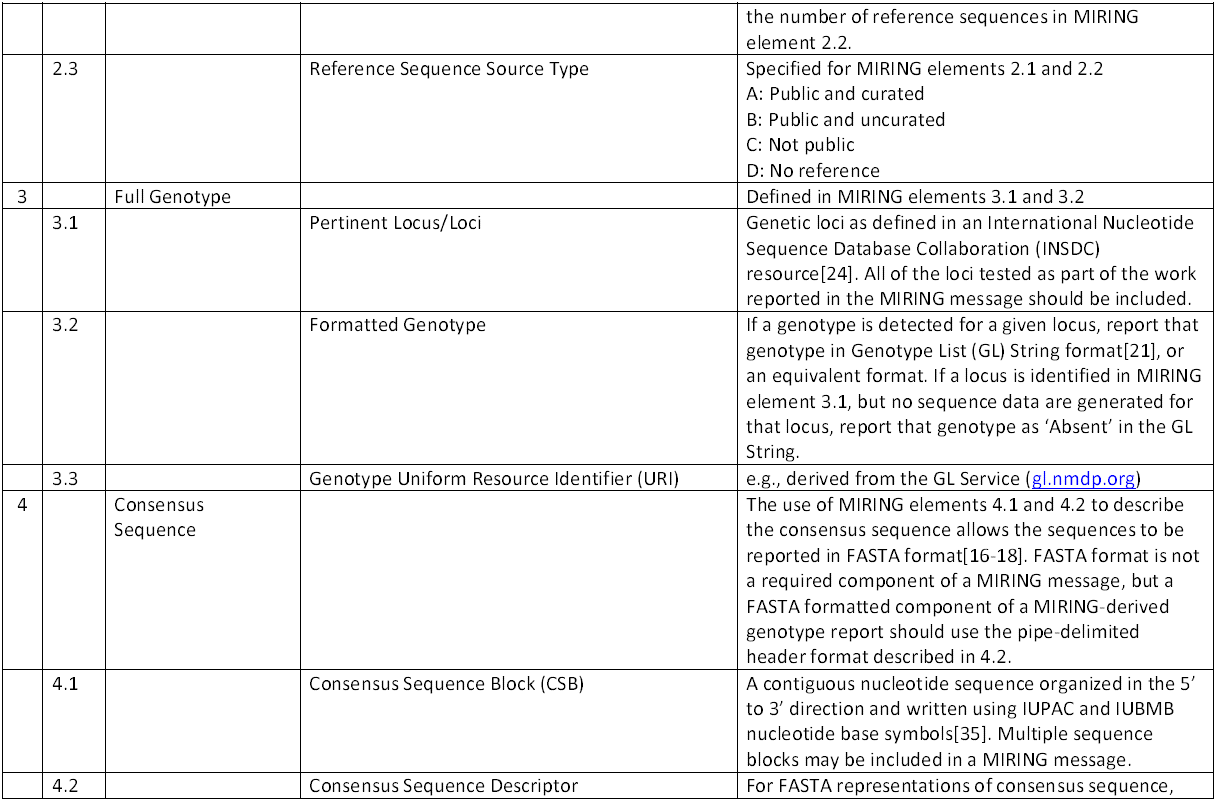
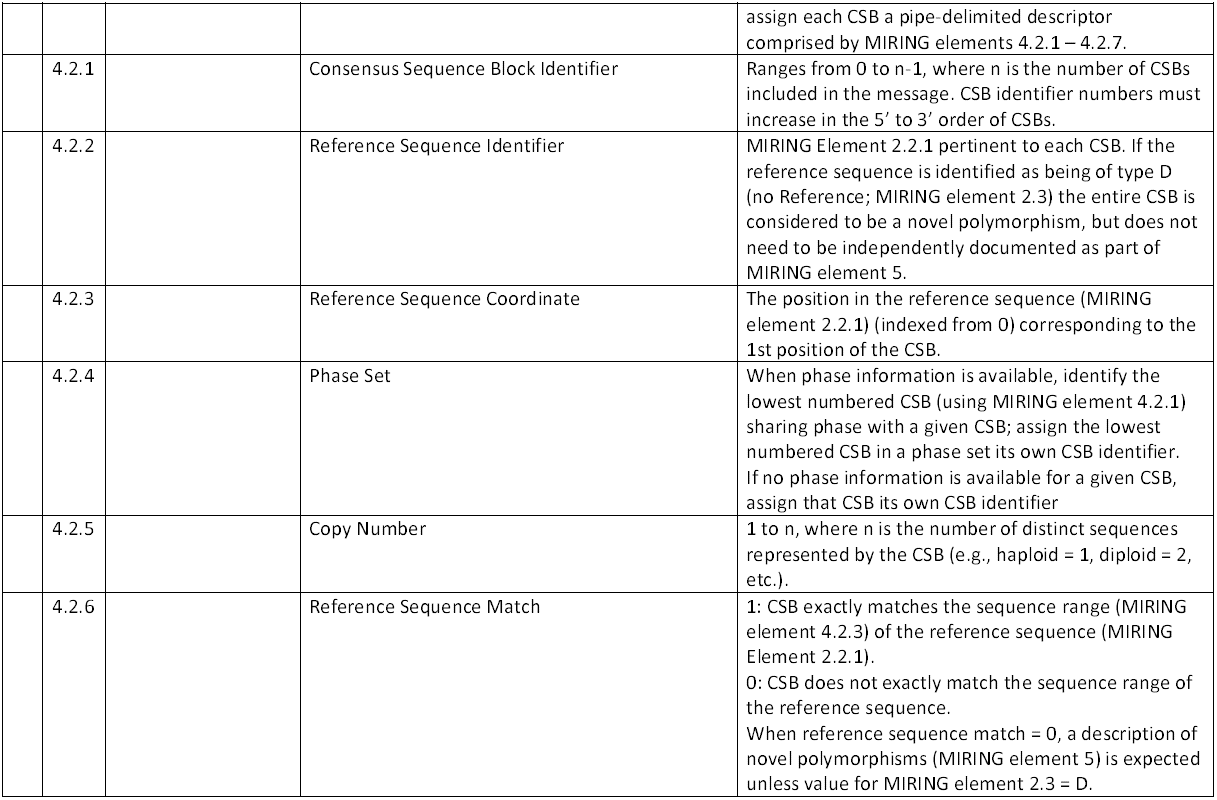
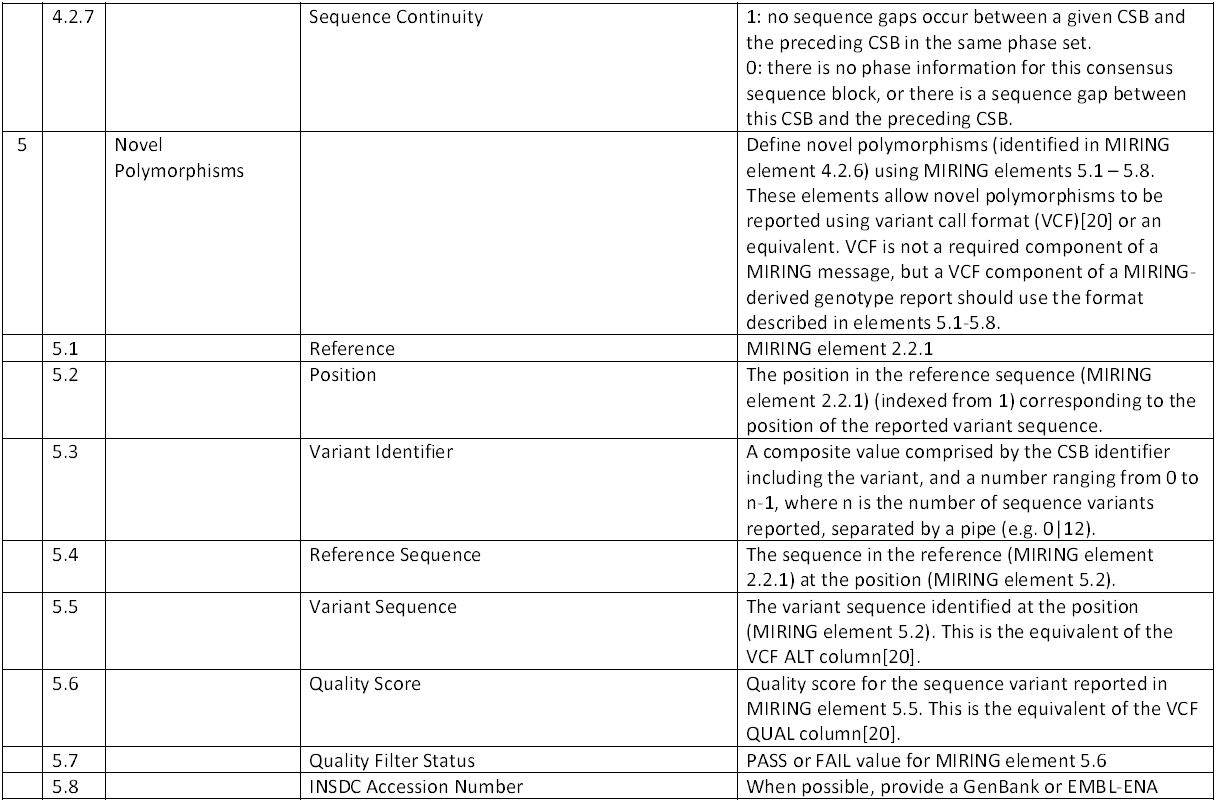
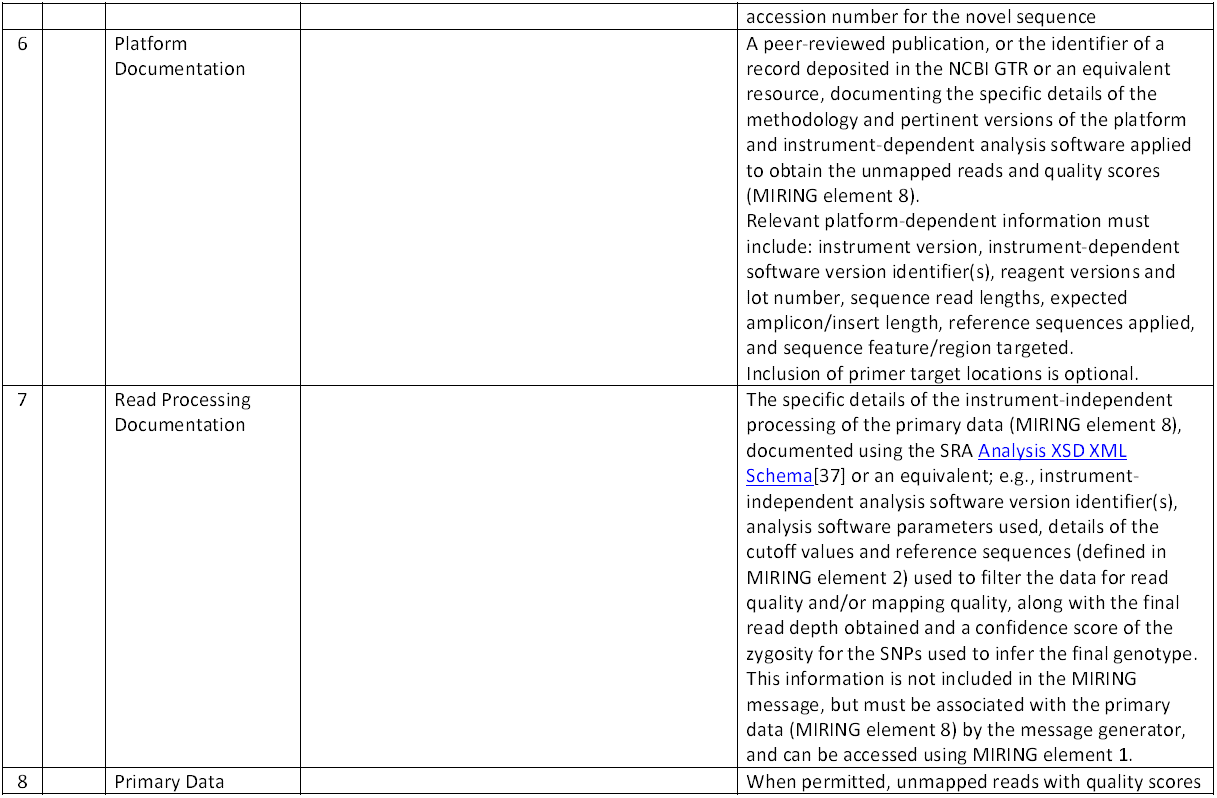
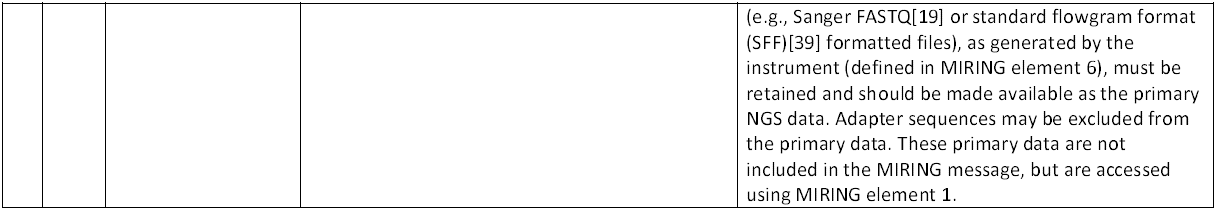
Definition of MIRING Elements and Formats

#### MIRING Element 1: Message Annotation

Each MIRING message must include a unique identifier that links the message contents to external information excluded from the MIRING message. For example, individual subject identifiers protected by the U.S. Health Insurance Portability and Accountability Act of 1996 (HIPAA)[26], the Canadian Personal Information Protection and Electronic Documents Act (PIPEDA)[27], and EU Directive 95/46/EC[28] are excluded from MIRING messages, and should be reported and transmitted outside the scope of the message. MIRING message annotation must allow unambiguous identification of the organization that generated the message, as well as the unambiguous identification of any MIRING message generated by that organization. For example, organizations can be identified unambiguously using an International Organization for Standardization (ISO)/International Electrotechnical Commission (IEC) standard 6523 organization identifier (OID)[29, 30]. MIRING message annotation must also include contact information for the organization that generated the message, along with references to the location and availability of platform (MIRING element 6) and read processing (MIRING element 7) documentation, and the primary data (MIRING element 8) from which the MIRING message was generated.

#### MIRING Element 2: Reference Context

Comparison to specific well-characterized and annotated reference sequences is crucial for NGS genotyping of *HLA* and *KIR* genes. Reference sequences may be used for the mapping and processing of reads, as well as for the determination of a genotype based on a consensus sequence. These sequences are stored in a variety of databases, including the Genome Reference Consortium (GRC)[31] and IMGT/HLA and IPD-KIR Databases [23]. Clear identification both of the database versions and of the individual sequences applied in an NGS genotyping must be included in each MIRING message. For example, GRCh38.p4 or IMGT/HLA Database release 3.21.1 describe the current versions of the GRC human genome and IMGT/HLA Databases, respectively; GL000251.2. and HLA00001 are the accession numbers for the GRCh38.p4 alternate reference locus number 2, (derived from the COX cell line) and IMGT/HLA Database *HLA-A*01:01:01:01* allele, respectively.

Any reference database or individual reference sequence used to generate a NGS genotype should be documented as part of MIRING element 2. To allow the assessment of the confidence in the genotyping, this documentation should indicate whether or not the database is public, and if a public database is curated. If no reference database is used, this should also be indicated. For instances when either previously unexplored gene features (e.g., *HLA-DRB5* introns) or a genomic region that is unrepresented in any genomic alignment (e.g., *DR1* or *DR8* haplotypes of the *HLA-DRB* region[32]) is being sequenced, the absence of available reference sequence at the time of the genotyping should be noted.

#### MIRING Element 3: Full Genotype

Some NGS methods provide phased, full-gene resolution data. However, many NGS genotyping approaches do not return such results, and genotyping ambiguity [7] is not resolved. In order to evaluate genotyping results across specimens, NGS instruments, genotyping and analysis methods, the complete set of alleles and genotype combinations that are possible for a given set of sequence data using a given reference sequence database must be provided.

GL String[22] format can be used to describe the genotype at each locus, including genotyping ambiguity and known allelic phase between loci. A “best guess,” estimate or imputation of an unambiguous genotype should <u>not</u> be included in a MIRING message. When available, a reference to an external uniform resource identifier (URI) for the GL String should be included as well.

In addition, genes that were specifically targeted, but yielded no sequence data should be explicitly identified. Due to structural variation among the *HLA*[32] and *KIR* loci[33–35], genes that are present in some individuals may be completely absent in others. For example, individuals homozygous for *HLA-DRB1*01* alleles have no *HLA-DRB3, HLA-DRB4* or *HLA-DRB5* genes[32], and individuals homozygous for the *KIR* A haplotype have no *KIR2DS1, KIR2DS3/5* or *KIR2DL5A* genes[33]. When a gene could have been detected by a given NGS instrument, but no sequence for that gene is generated for a subject, the locus in question should be identified in the MIRING message, with the genotype reported as “Absent” for that locus.

#### MIRING Element 4: Consensus Sequence

Depending on the NGS approach applied, consensus sequence for an individual reported allele may be generated as a single, gene-length consensus sequence block (CSB) (e.g., resulting from *de novo* assembly), or as shorter phased or unphased CSBs (e.g., corresponding to individual exons). These sequences should be written using the single-letter symbols for nucleotide bases and incompletely specified bases defined by the International Union of Pure and Applied Chemistry (IUPAC) and International Union of Biochemistry and Molecular Biology (IUBMB)[36]. Two CSBs would be reported for heterozygous individuals where complete phase was known for a given gene.

Each CSB should be accompanied by MIRING elements 4.2.1–4.2.7, which identify the CSB, the reference sequence (as defined in MIRING element 2) to which it has been aligned along with its position and identity to that reference sequence, or the absence of a reference sequence, any phase and continuity between CSBs, and the inferred copy number for each CSB. For diploid loci, the copy number values for homologous CSBs should sum to 2; however, due to copy number variation of some *HLA* and *KIR* genes[32, 33], some individuals are truly haploid for a given gene (copy number of 1), while others may have more than two copies of a given gene (e.g., copy number of 3 or 4). For example, some individuals have four copies of the *KIR2DS3* and *KIR2DL5* genes[33].

A CSB can be described in FASTA format by including a header line that comprises MIRING elements 4.2.1–4.2.7, formatted as defined in Table 1 and illustrated in Figure 1. Although this header format should be used for FASTA presentation of CSBs in a genotyping report, MIRING elements 4.2.1 – 4.2.7 can be recorded separately and differently within a MIRING message.

**Figure 1.**
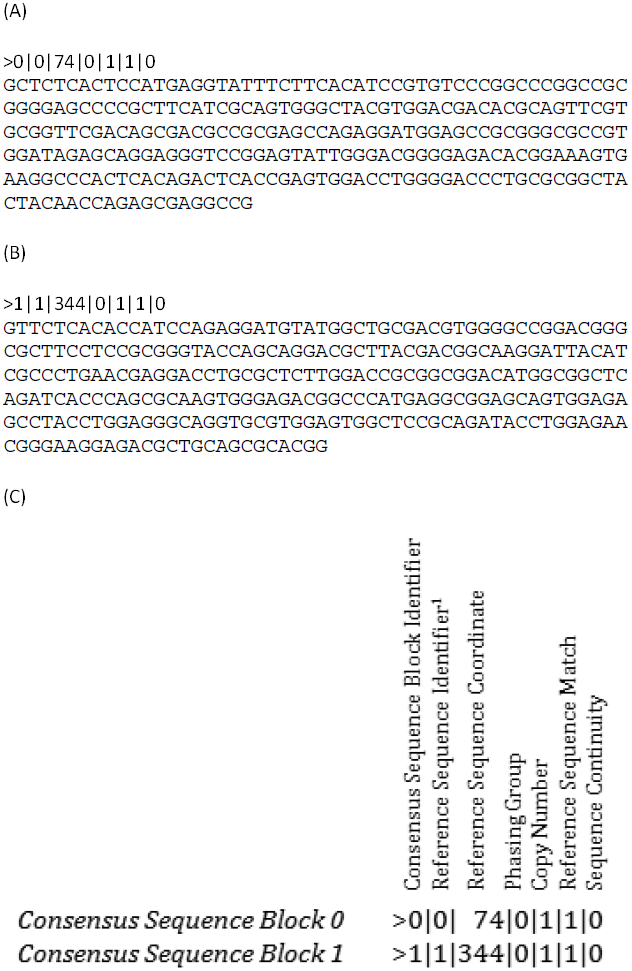
FASTA Consensus Sequence Blocks with MIRING Headers. Consensus sequences in MIRING messages are arranged in consensus sequence blocks (CSBs). CSBs are equivalent to FASTA formatted sequences that use a MIRING-specific descriptor as the header line. CSBs representing phased exon 2 and exon 3 sequences for an *HLA-A* allele are shown in A and 1B. A MIRING CSB descriptor consists of seven fields of information, delimited by pipes (|) as shown in C. Each CSB is identified with a unique index, beginning from 0, in the Consensus Sequence Block Identifier field. CSB identifiers must increase in the 5’ to 3’ direction of CSBs. Each reference sequence is identified with a unique index, beginning from 0, in the Reference Sequence Identifier field. Each index is defined in MIRING element 2. The Reference Sequence Coordinate field identifies the first position of the pertinent CSB in the pertinent reference sequence. The Phasing Group field identifies CSBs between which phase is known. Each set of phased CSBs will be identified with the same index, beginning from 0. The Copy Number field identifies the number of distinct sequences represented by each CSB (e.g., 1 = haploid, 2 = diploid, etc.). The Reference Sequence Match field identifies CSBs that exactly match the sequence range of the pertinent reference sequence (value = 1) or that do not match the sequence range of the reference sequence (value = 0). When phase is indicated for CSBs, the Sequence Continuity field indicates if there are gaps (of any size) between those phased CSBs. A value of 1 in this field indicates that there are no gaps between the pertinent CSB and the most immediately 5’ CSB in phase with that CSB. A value of 0 in this field indicates a sequence gap between the pertinent CSB and the most immediately 5’ CSB in phase, or that no phase information is available for the pertinent CSB. (A) CSB 0 and header, (B) CSB 1 and header, (C) Guide to interpreting the MIRING headers. 1: Reference sequence for CSB 0 is IMGT/HLA Database release version 3.21.1 A_nuc.fasta HLA00005 HLA-A*02:01:01:01. Reference sequence for CSB 1 is IMGT/HLA Database release version 3.21.1 A_nuc.fasta HLA10254 HLA-A*66:01:02.

#### MIRING Element 5: Novel Sequence Polymorphisms

The *HLA* and *KIR* genes are highly polymorphic, and the number of alleles reported to public databases is expected to increase dramatically with the use of NGS genotyping[37]. In the context of a MIRING message, novel polymorphism includes nucleotide sequence variants not yet present in a curated, public reference sequence database (e.g., the IMGT/HLA Database). The explicit identification of novel polymorphisms is an important element of both clinical and research genotyping, and must be documented in a MIRING message. Depending on how a MIRING message is generated, a CSB (as defined in MIRING element 4) representing novel sequence variants may have been submitted to a non-curated public reference sequence database [e.g., GenBank or the European Molecular Biology Laboratory (EMBL) European Nucleotide Archive (ENA)] as a novel sequence.

Novel sequence variants can be described through the use of MIRING elements 5.1–5.8, which define the reference sequence, variant position, variant sequence, quality score and quality filter status, and (if available) a GenBank or EMBL-ENA accession number for the novel sequence (e.g. L28096). This accession number can be included in the GL String (MIRING element 4.2.2, e.g. *HLA-DRB1\**L28096).

Use of MIRING elements 5.1–5.8 is sufficient to describe novel sequence variants in VCF as part of a genotyping report. A GenBank or EMBL-ENA accession number can be linked to the variant identifier (MIRING element 5.3) in the VCF meta-data. However, actual VCF is not a required component of a MIRING message, and MIRING elements 5.1–5.8 can be recorded separately and differently within a MIRING message.

#### MIRING Element 6: Platform Documentation

The specifics of each instrument, methodological approach (e.g., whole-genome sequencing, target enrichment, targeted amplicon sequencing) and reagent set applied to generate the primary read data (MIRING element 8) upon which the genotyping result is based should be documented. This documentation can take the form of a citation to a peer-reviewed publication or a reference to a structured documentation of the instrument and methodology in a publically accessible resource (e.g., the NCBI’s Genetic Testing Registry (GTR)). This documentation is not included in the MIRING message, but the reference to the resource must be included in MIRING element 1.

#### MIRING Element 7: Read Processing Documentation

After the primary read data (MIRING element 8) have been generated by the NGS instrument, they may be scrutinized for quality and length, modified or subjected to various bioinformatics filters before allele calls are made and a genotype result is generated. To enable the replication of the genotyping result and the evaluation of the bioinformatics process itself, the software and version used for both the genotyping and the processing steps applied must be documented. While these read processing steps are idiosyncratic to the combination of NGS components that have been applied in the genotyping effort, they can be accommodated in the NCBI’s Sequence Read Archive (SRA) Analysis XSD XML Schema[38, 39]. Each MIRING message should reference a report, based on the Analysis XSD or an equivalent, describing each program or script applied in the processing of reads, the order in which they were applied, the software versions and the pertinent parameters used. Where possible, this report should be associated with the primary read data (MIRING element 8), or made available by the MIRING message generator. This report should not be included in the MIRING message, but a reference to the location of the read processing documentation, or instructions for obtaining access to that documentation, should be included in MIRING element 1.

#### MIRING Element 8: Primary Data

The reads generated by the NGS instrument applied for the typing should be made available for re-analysis, either via deposition in a public database [e.g., the SRA, NCBI’s Genotype and Phenotype database (dbGAP) or an equivalent] or directly from the data generators. Primary data should take the form of unmapped reads with quality scores (e.g. Sanger FASTQ, SFF[40], or unmapped BAM including quality scores). The primary data are not included in the MIRING message, but a reference to the location of the primary data, or instructions for obtaining access to those data, should be included in MIRING element 1.

Because each MIRING message is assigned a unique identifier (MIRING element 1), it is possible for a MIRING message generator to produce multiple distinct messages for a single specimen from one set of primary data. Depending on read processing parameters applied and references used, each message may use a different subset of reads. Therefore, the MIRING message generator must maintain an archive for a given set of primary data, which identifies the reads pertinent to each MIRING message. When the primary data are publically available (as defined as part of MIRING element 1), the MIRING message generator must make this information available as well.

## 3. Strengths and Limitations of MIRING

MIRING represents a bold departure from previous methods of reporting *HLA* and *KIR* genotyping results. Previously, genotyping results have been maintained as static entities constrained by the existing references and nomenclature, with insufficient reference context and sequence information provided to foster genotype reassessment. By contrast, MIRING elements 3–5 are dynamic in that they can change with the reference context (MIRING element 2) - for instance when the database is updated. By providing access to the primary read data, and by including the consensus sequence in the MIRING message, the genotype and novel polymorphism information in a MIRING message can be updated with each reference allele sequence database revision.

The MIRING checklist identifies the minimum information needed to provide adequate documentation of an NGS *HLA* and *KIR* genotype. This documentation should be sufficient to reproduce or permit the reinterpretation of the reported genotype from the primary read data, and to allow informed comparisons of genotyping results for the same subject generated using different NGS genotyping instruments, reference sequences and analytical software. In cases when such genotyping results differ, use of MIRING messages to report those genotypes should facilitate the rapid identification of the sources of such discrepancies.

Many pieces of information pertinent to a genotyping experiment are not included in MIRING messages. Protected subject identifiers, phenotypic and demographic subject details, specimen details (e.g., preparation, quantification), specifics of the activity for which the genotyping effort was undertaken, descriptions and selection criteria for the loci genotyped, interpretations of the genotyping result (e.g., a ‘best call’ for ambiguous genotypes, or identifying a donor as a match to a patient), and funding sources for the genotyping are excluded from MIRING messages. Although this information can be associated with MIRING messages as part of a larger message if required, MIRING’s main purpose is to document those elements of a genotyping experiment that foster the archival utility of the genotyping result.

The MIRING checklist has been developed via community consensus, in order to meet the data management and exchange needs of the histocompatibility and immunogenetics community. Much of the information included in a MIRING message will not be pertinent to all current *HLA* and *KIR* genotyping use cases (e.g., clinical care, basic research, instrument validation, software development), but all such use cases can be met using the same MIRING message. Applications that parse MIRING messages and provide information tailored to each use case will make use of the same message. Given the rapid exploration of NGS technologies and methodologies, use of MIRING messages will allow the transparent evaluation of different instruments and genotyping approaches, encouraging improvement and standardization of all NGS methods.

The elements of the MIRING checklist allow MIRING messages to pertain to a single locus or multiple loci, but not to multiple subjects. Data for discrete subjects are reported in distinct MIRING messages, and multiple messages can be generated for each subject. Proper management of MIRING message identifiers then becomes an essential part of the MIRING messaging system. A central repository for MIRING messages will greatly facilitate and simplify the exchange of genotyping results among centers and researchers.

As illustrated in Figure 2, the elements of a MIRING message can be divided into distinct categories. MIRING elements 6–8 are specific to NGS methodologies, but are not included in the MIRING message. Therefore, MIRING messaging could be extended to include other genotyping methodologies (e.g., SBT, SSOP or SSP) by broadening the scope of information reported in elements 6–8 to document the methodologies, software, and raw data pertinent to these other methodologies. However, the structure or content of MIRING elements 1–5 would not change. Only the details of the MIRING message would change; for example, CSBs would decrease in length for SSOP- and SSP-based MIRING messages, to accommodate the shorter length of the hybridized sequences[41], and ambiguity in the full genotypes reported would increase. CSB copy number values would change from 1 (haploid) to 2 (diploid) for most Sanger sequencing-based MIRING messages (unless sequencing was applied to isolated chromosomes, or on the basis of group-specific amplification[42]).

**Figure 2.**
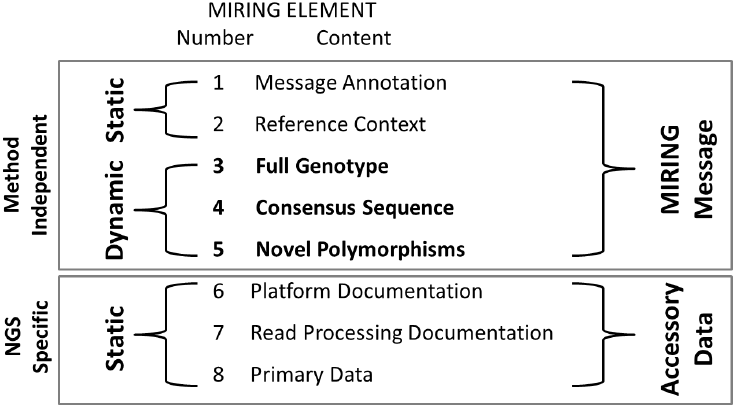
MIRING Checklist Categories. The eight elements of the MIRING checklist are either included in a MIRING message or exist as accessory data that are referenced in the MIRING message. MIRING message elements 1 and 2 and 6–8 are static in that they pertain to events that occurred when the genotyping instrument was applied to generate the primary read data. MIRING elements 3–5 are dynamic in that they can change if MIRING element 2, changes (e.g., a future release of the IPD-KIR database may result in a change to the ambiguity level of a genotype). MIRING elements 6–8 are also specific to NGS platforms. Valid MIRING messages could be generated on the basis of Sanger sequence-based typing (SBT), sequence-specific oligonucleotide probe (SSOP) and priming (SSP) methods, requiring changes to the content of MIRING elements 6–8 alone.

Although, the MIRING checklist was developed with *HLA* and *KIR* genotypes in mind, it can be applied to report genotype data for any highly-polymorphic genetic system. In addition, MIRING is sufficiently flexible that it can accommodate future developments in sequencing technology. As the cost of generating high-quality genomic information decreases, the need to report and exchange these data in a straightforward and reproducible manner will increase. As genomic data accumulate, specific genes, haplotype-blocks and chromosomal regions will be revealed as medically relevant, and their polymorphism can be documented and reported via a MIRING message.

Examples of Histoimmunogenetics Markup Language version 1.0 messages that comply with MIRING standards and principles are included in the paper by Milius et al. [43] included in this issue.

## 4. Conclusions

MIRING messages foster the portability of *HLA* and *KIR* genotype data in a standard format, allowing the dynamic re-analysis of these medically important results in the context of continual genomic discovery. The data recorded in a MIRING message are essential for the systematic traceability of a NGS genotyping result; this traceability is critical for reproducible research and the meaningful archiving of modern genotyping results. The MIRING checklist is sufficiently broad in scope that genotyping results generated using NGS technologies, older genotyping technologies and future methods can be accommodated in a MIRING message. Reporting of NGS genotype data as MIRING messages promotes transparency across varying applications of NGS components, facilitating comparison, and therefore the improvement and ongoing development of NGS technology. Finally, the widespread application of the MIRING checklist for exchanging NGS genotyping results will allow the leveraging of public data resources for meta-analysis, study replication and new discovery. More information about MIRING can be found online at miring.immunogenomics.org.

## Acknowledgements

This work was supported by National Institutes of Health (NIH) grants U01AI067068 (SJM and JAH) and U01AI090905 (PRP and PJN), awarded by the National Institute of Allergy and Infectious Disease (NIAID), and R01GM109030 (SJM and JAH), awarded by the National Institute of General Medical Sciences (NIGMS), and by Office of Naval Research (ONR) grant N00014-08-1-1207 (MM and RPM). The content presented is solely the responsibility of the authors and does not necessarily represent the official views of the NIH, NIAID, NIGMS, ONR, Department of Defense or United States Government. We thank all of the participants in the INGSDC for their input and useful discussions regarding the development of MIRING.

